# RASCL: Rapid Assessment Of SARS-CoV-2 Clades Through Molecular Sequence Analysis

**DOI:** 10.1101/2022.01.15.476448

**Authors:** Alexander G Lucaci, Jordan D Zehr, Stephen D Shank, Dave Bouvier, Han Mei, Anton Nekrutenko, Darren P Martin, Sergei L Kosakovsky Pond

## Abstract

An important component of efforts to manage the ongoing COVID19 pandemic is the <u>R</u>apid <u>A</u>ssessment of how natural selection contributes to the emergence and proliferation of potentially dangerous <u>S</u>ARS-CoV-2 lineages and <u>CL</u>ades (RASCL). The RASCL pipeline enables continuous comparative phylogenetics-based selection analyses of rapidly growing clade-focused genome surveillance datasets, such as those produced following the initial detection of potentially dangerous variants. From such datasets RASCL automatically generates down-sampled codon alignments of individual genes/ORFs containing contextualizing background reference sequences, analyzes these with a battery of selection tests, and outputs results as both machine readable JSON files, and interactive notebook-based visualizations.

**Availability:** RASCL is available from a dedicated repository at https://github.com/veg/RASCL and as a Galaxy workflow https://usegalaxy.eu/u/hyphy/w/rascl. Existing clade/variant analysis results are available here: https://observablehq.com/@aglucaci/rascl.

**Contact:** Dr. Sergei L Kosakovsky Pond.

**Supplementary information:** N/A

---

Rapid characterization and assessment of the clade-specific molecular features of individual persistent or rapidly expanding SARS-CoV-2 lineages has become an important component of efforts to monitor and manage the COVID19 pandemic. Analyses of natural selection have been broadly incorporated into such assessments as a primary tool for inferring the selective processes under which novel SARS-CoV-2 variants evolve (Tegally *et al*., 2021, Faria *et al*., 2021, D. Martin *et al*., 2021, MacLean et al., 2021). Ongoing monitoring of emergent variants of interest (VOI) or concern (VOC) can detect potentially adaptive mutations before they rise to high frequency, and help establish the relationships between individual mutations and key viral characteristics including pathogenicity, transmissibility, and drug resistance (Hamed *et al*., 2021, Young *et al*., 2021, Luchsinger *et al*., 2021, Abdool *et*.*al*., 2021, Cyrus Maher *et al*., 2021). Molecular patterns of ongoing selection that are evident within sequences sampled from particular VOI or VOC clades may also reveal the sub-lineages within these clades that carry potentially fitness-enhancing mutations and which are therefore most likely to drive future viral transmission (Rambaut *et al*., 2020).

Here, we present RASCL (**<u>R</u>**apid **<u>A</u>**ssessment of **<u>S</u>**ARS-CoV-2 **<u>CL</u>**ades), an analytic pipeline designed to investigate the nature and extent of selective forces acting on viral genes in SARS-CoV-2 sequences through comparative phylogenetic analyses (Figure 1A). RASCL is implemented as an easy-to-use, standalone pipeline and as a web application, integrated in the Galaxy framework and available for use on powerful public computing infrastructure (Afgan *et al*., 2018).

**Figure 1.**
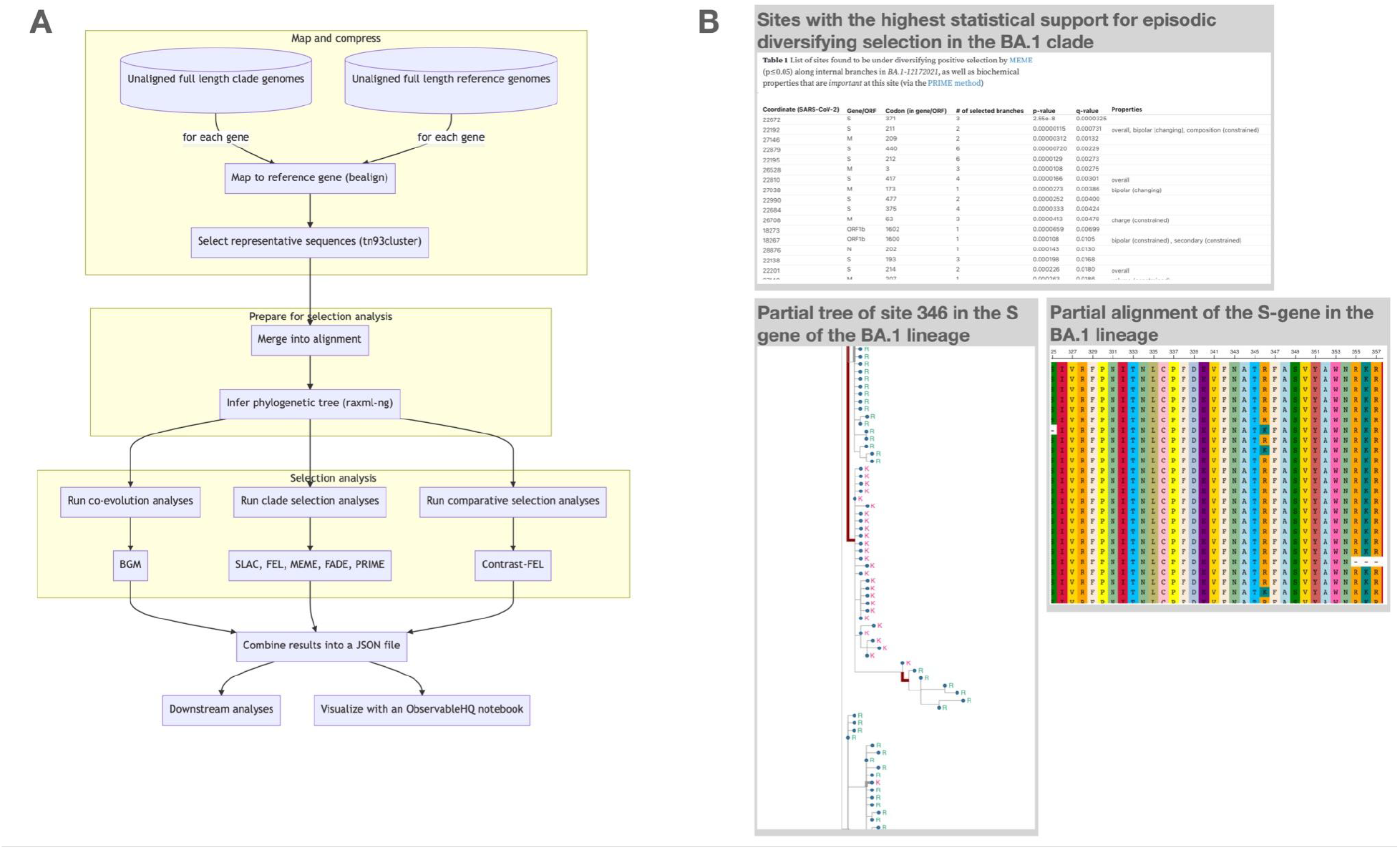
(A) A flowchart diagram of the main analytic engine of RASCL. (B) Examples of the ObservableHQ visualization notebook elements for the main Omicron clade (BA.1).

The RASCL pipeline takes as input (i) a “query” dataset comprising a single FASTA file containing unaligned SARS-CoV-2 full or partial genomes belonging to a clade of interest (e.g., all sequences from the PANGO lineage, B.1.617.2) and (ii) a generic “background” dataset that might comprise, for example, a set of sequences that are representative of global SARS-CoV-2 genomic diversity assembled from ViPR (Pickett *et al*., 2012). It is not necessary to remove sequences in the query dataset from the reference dataset -- the pipeline will do this automatically. The choice of “query” and “background” datasets is analysis-specific. For example, if another clade of interest is provided as background it is possible to identify sites that are evolving differently between two clades directly. Other sensible choices of query sequences might be: sequences from a specific country/region, or sequences sampled during a particular time period. Following the disassembly of whole genome datasets into individual coding sequences (based on the NCBI SARS-CoV-2 reference annotation), the gene datasets (each containing a set of query and background sequences) are processed in parallel.

Using complete linkage distance clustering implemented in the TN93 package (https://github.com/veg/tn93), RASCL subsamples from available sequences while attempting to maintain genomic diversity; the clustering threshold distance is chosen automatically to include no more than a user-specified number of genomes (e.g., 300). A combined (query and background) alignment is created with only the sequences that are divergent enough to be useful for subsequent selection analyses being retained from the background dataset. Inference of a maximum likelihood phylogenetic tree (RAxML-NG, Kozlov *et al*., 2019, or IQ-TREE, Nguyen *et al*., 2015) is performed on the combined dataset and the query and background branches of this tree are labeled. Selection analyses are then performed with state of the art molecular evolution models implemented in HyPhy (Pond *et al*., 2020).

1. SLAC: performs substitution mapping (Pond and Frost, 2005)
2. BGM: identifies groups of sites that are apparently co-evolving (Poon *et al*., 2008)
3. FEL: locates codon sites with evidence of pervasive positive diversifying or negative selection (Pond and Frost, 2005),
4. MEME: locates codon sites with evidence of episodic positive diversifying selection, (Murrell *et al*., 2012)
5. BUSTEDS: tests for gene-wide episodic selection (Wisotsky *et al*., 2020)
6. RELAX: compare gene-wide selection pressure between the query clade and background sequences (Wertheim *et al*., 2015),
7. CFEL: comparison site-by-site selection pressure between query and background sequences (Pond *et al*., 2021).
8. FADE: identify amino-acid sites with evidence of directional selection (Pond et.al., 2008)

To mitigate the potentially confounding influences of within-host evolution and sequencing errors, these analyses are performed only on internal branches of the phylogenetic tree (Lorenzo-Redondo *et al*., 2016). Results are combined into two machine readable JSON files (“Summary” and “Annotation”) that are used for web processing. A feature-rich interactive notebook in ObservableHQ (Perkel 2021, https://observablehq.com/@aglucaci/rascl) is used to visualize and summarize RASCL results (Figure 1B)

RASCL is currently available in two distributions:(i) through a web interface via the Galaxy Project as a workflow (https://usegalaxy.eu/u/hyphy/w/rascl); and (ii) as a standalone pipeline via a dedicated GitHub (https://github.com/veg/RASCL) repository. For the web application implementation, the alignment, tree and analysis results are stored and made web-accessible via the Galaxy platform. Results are visualized with an interactive notebook hosted on ObservableHQ (Figure 1B; Perkel 2021) that includes an alignment viewer, a visualization of individual codons/amino acid states at user-selected sites mapped onto the tips of a phylogenetic tree, and detailed tabulated information on analysis results for individual genes and codon-sites.

RASCL has been used to characterize the role of natural selection in the emergence of the Beta (Tegally *et al*., 2021), Gamma (Faria *et al*., 2021), and Omicron (Moyo *et al*., 2021) VOC lineages, and for identifying patterns of convergent evolution in N501Y SARS-CoV-2 lineages (Martin et al., 2021). Whenever future genomic surveillance efforts reveal new potentially problematic SARS-CoV-2 lineages, we anticipate that RASCL will be productively used to analyze these too. Finally, RASCL has been designed so that, with minimal modification, it can also be adapted to analyze any other viral pathogens for which sufficient sequencing data is available.

## Acknowledgements

We thank members of the Datamonkey/HyPhy and Galaxy teams for their assistance in the development of this application. DPM is funded by the Wellcome Trust (222574/Z/21/Z). This research was supported in part by grants R01 AI134384 (NIH/NIAID) and grant 2027196 (NSF/DBI,BIO) to AN and SLKP..

## References

1. Tegally, H., Wilkinson, E., Giovanetti, M. et al. Detection of a SARS-CoV-2 variant of concern in South Africa. Nature 592, 438–443 (2021). https://doi.org/10.1038/s41586-021-03402-9

2. Faria NR, Mellan TA, Whittaker C, Claro IM, Candido DDS, Mishra S, Crispim MAE, Sales FCS, Hawryluk I, McCrone JT, Hulswit RJG, Franco LAM, Ramundo MS, de Jesus JG, Andrade PS, Coletti TM, Ferreira GM, Silva CAM, Manuli ER, Pereira RHM, Peixoto PS, Kraemer MUG, Gaburo N Jr, Camilo CDC, Hoeltgebaum H, Souza WM, Rocha EC, de Souza LM, de Pinho MC, Araujo LJT, Malta FSV, de Lima AB, Silva JDP, Zauli DAG, Ferreira ACS, Schnekenberg RP, Laydon DJ, Walker PGT, Schlüter HM, Dos Santos Alp, Vidal MS, Del Caro VS, Filho RMF, Dos Santos HM, Aguiar RS, Proença-Modena Jl, Nelson B, Hay JA, Monod M, Miscouridou X, Coupland H, Sonabend R, Vollmer M, Gandy A, Prete CA Jr, Nascimento VH, Suchard MA, Bowden TA, Pond SLK, Wu CH, Ratmann O, Ferguson NM, Dye C, Loman NJ, Lemey P, Rambaut A, Fraiji NA, Carvalho Mdpss, Pybus OG, Flaxman S, Bhatt S, Sabino EC. Genomics and epidemiology of the P.1 SARS-CoV-2 lineage in Manaus, Brazil. Science. 2021 May 21;372(6544):815–821. doi: 10.1126/science.abh2644. Epub 2021 Apr 14. PMID: 33853970; PMCID: PMC8139423.

3. Elbe, S., and Buckland-Merrett, G. (2017) Data, disease and diplomacy: GISAID’s innovative contribution to global health. Global Challenges, 1:33–46. DOI:10.1002/gch2.1018 PMCID: 31565258

4. Pickett BE, Sadat EL, Zhang Y, Noronha JM, Squires RB, Hunt V, Liu M, Kumar S, Zaremba S, Gu Z, Zhou L, Larson CN, Dietrich J, Klem EB, Scheuermann RH. ViPR: an open bioinformatics database and analysis resource for virology research. Nucleic Acids Res. 2012 Jan;40(Database issue):D593–8. doi: 10.1093/nar/gkr859. Epub 2011 Oct 17. PMID: 22006842; PMCID: PMC3245011.

5. Sergei L Kosakovsky Pond, Art FY Poon, Ryan Velazquez, Steven Weaver, N Lance Hepler, Ben Murrell, Stephen D Shank, Brittany Rife Magalis, Dave Bouvier, Anton Nekrutenko, Sadie Wisotsky, Stephanie J Spielman, Simon DW Frost, Spencer V Muse (2020) HyPhy 2.5—A Customizable Platform for Evolutionary Hypothesis Testing Using Phylogenies. Molecular Biology and Evolution 37.1 (2020): 295–299.

6. Steven Weaver, Stephen D. Shank, Stephanie J. Spielman, Michael Li, Spencer V. Muse, Sergei L. Kosakovsky Pond Datamonkey 2.0: a modern web application for characterizing selective and other evolutionary processes Mol. Biol. Evol. 35(3):773–777

7. Tamura K, Nei M. Estimation of the number of nucleotide substitutions in the control region of mitochondrial DNA in humans and chimpanzees. Mol Biol Evol. 1993 May;10(3):512–26. doi: 10.1093/oxfordjournals.molbev.a040023. PMID: 8336541.

8. Alexey M Kozlov, Diego Darriba, Tomáš Flouri, Benoit Morel, Alexandros Stamatakis, RAxML-NG: a fast, scalable and user-friendly tool for maximum likelihood phylogenetic inference, Bioinformatics, Volume 35, Issue 21, 1 November 2019, Pages 4453–4455, https://doi.org/10.1093/bioinformatics/btz305

9. Sergei L. Kosakovsky Pond and Simon D. W. Frost (2005) Not So Different After All: A Comparison of Methods for Detecting Amino Acid Sites Under Selection Molecular Biology and Evolution 22(5): 1208–1222

10. Art F. Y. Poon, Fraser I. Lewis, Simon D. W. Frost, Sergei L. Kosakovsky Pond, Spidermonkey: rapid detection of co-evolving sites using Bayesian graphical models, Bioinformatics, Volume 24, Issue 17, 1 September 2008, Pages 1949–1950, https://doi.org/10.1093/bioinformatics/btn313

11. Ben Murrell, Joel O. Wertheim, Sasha Moola, Thomas Weighill, Konrad Scheffler and Sergei L. Kosakovsky Pond (2012) Detecting Individual Sites Subject to Episodic Diversifying Selection PLoS Genetics 8(7): e1002764

12. M. D. Smith, J. O. Wertheim, S. Weaver, B. Murrell, K. Scheffler and S. L. Kosakovsky Pond Less Is More: An Adaptive Branch-Site Random Effects Model for Efficient Detection of Episodic Diversifying Selection Molecular Biology and Evolution 32: 1342–1353

13. Wisotsky SR, Kosakovsky Pond SL, Shank SD, Muse SV. Synonymous Site-to-Site Substitution Rate Variation Dramatically Inflates False Positive Rates of Selection Analyses: Ignore at Your Own Peril. Mol Biol Evol. 2020 Aug 1;37(8):2430–2439. doi: 10.1093/molbev/msaa037. PMID: 32068869; PMCID: PMC7403620.

14. Wertheim JO, Murrell B, Smith MD, Kosakovsky Pond SL, Scheffler K. RELAX: detecting relaxed selection in a phylogenetic framework. Mol Biol Evol. 2015 Mar;32(3):820–32. doi: 10.1093/molbev/msu400. Epub 2014 Dec 23. PMID: 25540451; PMCID: PMC4327161.

15. Kosakovsky Pond SL, Wisotsky SR, Escalante A, Magalis BR, Weaver S. Contrast-FEL-A Test for Differences in Selective Pressures at Individual Sites among Clades and Sets of Branches. Mol Biol Evol. 2021 Mar 9;38(3):1184–1198. doi: 10.1093/molbev/msaa263. PMID: 33064823; PMCID: PMC7947784.

16. Hamed, S.M., Elkhatib, W.F., Khairalla, A.S. et al. Global dynamics of SARS-CoV-2 clades and their relation to COVID-19 epidemiology. Sci Rep 11, 8435 (2021). https://doi.org/10.1038/s41598-021-87713-x

17. Association of SARS-CoV-2 clades with clinical, inflammatory and virologic outcomes: An observational study. Young, Barnaby E et al. EBioMedicine, Volume 66, 103319

18. Rambaut, A., Loman, N., Pybus, O., Barclay, W., Barrett, J., Carabelli, A., … & Volz, E. (2020). Preliminary genomic characterisation of an emergent SARS-CoV-2 lineage in the UK defined by a novel set of spike mutations. Genom. Epidemiol.

19. Luchsinger LL, Hillyer CD. Vaccine efficacy probable against COVID-19 variants. Science. 2021 Mar 12;371(6534):1116. doi: 10.1126/science.abg9461. PMID: 33707257.

20. Abdool Karim SS, de Oliveira T. New SARS-CoV-2 Variants - Clinical, Public Health, and Vaccine Implications. N Engl J Med. 2021 May 13;384(19):1866–1868. doi: 10.1056/NEJMc2100362. Epub 2021 Mar 24. PMID: 33761203; PMCID: PMC8008749.

21. Perkel JM. Reactive, reproducible, collaborative: computational notebooks evolve. Nature. 2021 May;593(7857):156–157. doi: 10.1038/d41586-021-01174-w. PMID: 33941927.

22. Enis Afgan, Dannon Baker, Bérénice Batut, Marius van den Beek, Dave Bouvier, Martin Čech, John Chilton, Dave Clements, Nate Coraor, Björn A Grüning, Aysam Guerler, Jennifer Hillman-Jackson, Saskia Hiltemann, Vahid Jalili, Helena Rasche, Nicola Soranzo, Jeremy Goecks, James Taylor, Anton Nekrutenko, Daniel Blankenberg, The Galaxy platform for accessible, reproducible and collaborative biomedical analyses: 2018 update, Nucleic Acids Research, Volume 46, Issue W1, 2 July 2018, Pages W537–W544, https://doi.org/10.1093/nar/gky379

23. L.-T. Nguyen, H.A. Schmidt, A. von Haeseler, B.Q. Minh (2015) IQ-TREE: A fast and effective stochastic algorithm for estimating maximum likelihood phylogenies.. Mol. Biol. Evol., 32:268–274. https://doi.org/10.1093/molbev/msu300

24. Martin, D. P., Weaver, S., Tegally, H., San, J. E., Shank, S. D., Wilkinson, E., Lucaci, A. G., Giandhari, J., Naidoo, S., Pillay, Y., Singh, L., Lessells, R. J.,NGS-SA, COVID-19 Genomics UK (COG-UK), Gupta, R. K., Wertheim, J. O., Nekturenko, A., Murrell, B., Harkins, G. W., Lemey, P., … Kosakovsky Pond, S. L. (2021). The emergence and ongoing convergent evolution of the SARS-CoV-2 N501Y lineages. Cell, 184(20), 5189–5200.e7. https://doi.org/10.1016/j.cell.2021.09.003

25. MacLean, Oscar A., Spyros Lytras, Steven Weaver, Joshua B. Singer, Maciej F. Boni, Philippe Lemey, Sergei L. Kosakovsky Pond, and David L. Robertson. “Natural selection in the evolution of SARS-CoV-2 in bats created a generalist virus and highly capable human pathogen.” PLoS biology 19, no. 3 (2021): e3001115.

26. Predicting the mutational drivers of future SARS-CoV-2 variants of concern M. Cyrus Maher, Istvan Bartha, Steven Weaver, Julia di Iulio, Elena Ferri, Leah Soriaga, Florian A. Lempp, Brian L. Hie, Bryan Bryson, Bonnie Berger, David L. Robertson, Gyorgy Snell, Davide Corti, Herbert W. Virgin, Sergei L. Kosakovsky Pond, Amalio Telenti medRxiv 2021.06.21.21259286; doi: https://doi.org/10.1101/2021.06.21.21259286

27. Kosakovsky Pond, S. L., Poon, A. F., Leigh Brown, A. J., & Frost, S. D. (2008). A maximum likelihood method for detecting directional evolution in protein sequences and its application to influenza A virus. Molecular biology and evolution, 25(9), 1809–1824. https://doi.org/10.1093/molbev/msn123

28. Lorenzo-Redondo R, Fryer HR, Bedford T, Kim EY, Archer J, Pond SLK, Chung YS, Penugonda S, Chipman J, Fletcher CV, Schacker TW, Malim MH, Rambaut A, Haase AT, McLean AR, Wolinsky SM. Persistent HIV-1 replication maintains the tissue reservoir during therapy. Nature. 2016 Feb 4;530(7588):51–56. doi: 10.1038/nature16933. Epub 2016 Jan 27. PMID: 26814962; PMCID: PMC4865637.

29. Viana R, Moyo S, Amoako DG, Tegally H, Scheepers C, Althaus CL, Anyaneji UJ, Phillip A Bester Maciej F Boni, Mohammed Chand, Wonderful T Choga, Rachel Colquhoun, Michaela Davids, Koen Deforche, Deelan Doolabh, Susan Engelbrecht, Josie Everatt, Jennifer Giandhari, Marta Giovanetti Diana Hardie Verity Hill, Nei-Yuan Hsiao, Arash Iranzadeh, Arshad Ismail, Charity Joseph, Rageema Joseph, Legodile Koopile, Sergei L Kosakovsky Pond, Moritz UG Kraemer, Lesego Kuate-Lere, Oluwakemi Laguda-Akingba Onalethatha Lesetedi-Mafoko, Lessells RJ, Shahin Lockman Alexander G Lucaci, Arisha Maharaj, Boitshoko Mahlangu, Tongai Maponga, Kamela Mahlakwane Zinhle Makatini, Gert Marais Dorcas Maruapula, Kereng Masupu, Mogomotsi Matshaba, Simnikiwe Mayaphi, Nokuzola Mbhele, Mpaphi B Mbulawa, Adriano Mendes, Koleka Mlisana Anele Mnguni, Thabo Mohale, Monika Moir, Kgomotso Moruisi, Mosepele Mosepele Gerald Motsatsi, Modisa S Motswaledi Thongbotho Mphoyakgosi, Nokukhanya Msomi, Peter N Mwangi Yeshnee Naidoo, Noxolo Ntuli, Martin Nyaga Lucier Olubayo Pillay S, Botshelo Radibe, Ramphal Y, Ramphal U, San JE, Lesley Scott, Roger Shapiro Lavanya Singh, Pamela Smith-Lawrence, Wendy Stevens, Amy Strydom, Kathleen Subramoney, Naume Tebeila, Derek Tshiabuila, Joseph Tsui, Stephanie van Wyk, Steven Weaver, Constantinos K Wibmer, Eduan Wilkinson, Nicole Wolter, Alexander E Zarebski, Boitumelo Zuze, Dominique Goedhals Wolfgang Preiser, Florette Treurnicht, Marietje Venter, Carolyn Williamson. Oliver G Pybus, Jinal Bhiman. Allison Glass, Martin DP, Rambaut A, Gaseitsiwe S, von Gottberg A, de Oliveira T. Rapid epidemic expansion of the SARS-CoV-2 Omicron variant in southern Africa medRxiv,MEDRXIV-2021-268028v1-deOliveira: (2021).

30. Tegally H, Wilkinson E, Giovanetti M, et al. Detection of a SARS-CoV-2 variant of concern in South Africa. Nature. 2021 Apr;592(7854):438-443. DOI: 10.1038/s41586-021-03402-9. PMID: 33690265.

31. Faria Nuno R., Mellan Thomas A., Whittaker Charles, Claro Ingra M., Candido Darlan da S., Mishra Swapnil, Crispim Myuki A.E., et al. “Genomics and Epidemiology of the P.1 SARS-CoV-2 Lineage in Manaus, Brazil.” Science 372, no. 6544 (May 21, 2021): 815–21. https://doi.org/10.1126/science.abh2644.

